# Active diffusion and advection in the *Drosophila* ooplasm result from the interplay of the actin and microtubule cytoskeletons

**DOI:** 10.1101/098590

**Authors:** Maik Drechsler, Fabio Giavazzi, Roberto Cerbino, Isabel M. Palacios

## Abstract

Transport in cells occurs via a delicate interplay of passive and active processes, including diffusion, directed transport and advection. Despite progresses in super-resolution microscopy, discriminating and quantifying these processes is a challenge, requiring tracking of rapidly moving, sub-diffraction objects in a crowded, noisy environment. Here we use Differential Dynamic Microscopy with different contrast mechanisms to provide a thorough characterization of the dynamics in the *Drosophila* oocyte. We study the movement of vesicles and the elusive motion of a cytoplasmic F-actin mesh, a known regulator of cytoplasmic flows. We find that cytoplasmic motility constitutes a combination of directed motion and random diffusion. While advection is mainly attributed to microtubules, we find that active diffusion is driven by the actin cytoskeleton, although it is also enhanced by the flow. We also find that an important dynamic link exists between vesicles and cytoplasmic F-actin motion, as recently suggested in mouse oocytes.

## INTRODUCTION

The spatial distribution and organization of cytoplasmic content, like proteins or protein complexes, nucleic acids, and whole organelles require the combined action of passive and active biophysical processes. In fact, thermal-based diffusion is not sufficiently fast and effective in redistributing large organelles, like vesicles, within the cell^1^. This is due to their large size, which makes their diffusion coefficient small, as well as to the typically crowded and viscous environment that is found inside cells. Active transport mechanisms mitigate the ineffectiveness of thermal diffusion. Molecular motor proteins carry attached cargos (such as organelles and vesicles) along cytoskeletal filaments, which act as tracks for directed transport across the cell^2^. In addition, in larger cells, it is likely that the transport of such cargoes causes a large-scale net flow, known as cytoplasmic streaming^3,4^. As a result of the viscous drag, caused by a translocating motor, streaming leads to a circulation of the cytoplasm and its efficient remixing^5,6^. Finally, recent studies demonstrate that ATP-dependent processes are responsible for the presence of random force fluctuations within the cytoplasm, whose effects lead to the displacement of tracer particles in a diffusive-like manner - named active diffusion - that is more efficient than thermal-based diffusion^7–11^. Understanding the details of the subtle interplay between all these processes is a demanding task, due to the many time and length-scales and the multiple molecular pathways involved.

In this work, we investigate the interactions between different mechanisms of motion in the *Drosophila* oocyte. *Drosophila* oogenesis is well studied from a genetic point of view and this large cell can be probed with a variety of chemical treatments and microscopic tools. In the oocyte, microtubules and kinesin-1 are essential for both the transport of cargos and cytoplasmic streaming^12,13^. At mid-oogenesis (stage 9, st9), the topology and speed of cytoplasmic flows directly correlates with the topology of the microtubule cytoskeleton and the speed of kinesin, respectively^14^. In addition, a cytoplasmic network of actin filaments (F-actin) - known as the actin mesh - is present within the oocyte and acts as a negative regulator of the microtubule/kinesin-dependent flow^15^. However, much remains to be uncovered about the interplay between the actin mesh and the microtubule cytoskeleton in regulating the motion of material within the ooplasm. The recent discovery of a link between cytoplasmic actin and vesicle dynamics in mouse oocytes already suggests a close relationship between vesicle transport, motor activity and the actin cytoskeleton^16^. However, it remains unclear whether this observation represents a general feature, also present in other eukaryotic cells.

To gain insight on these issues we developed a methodology to simultaneously probe the dynamics of both, cytoplasmic F-actin and vesicles in wild type oocytes, as well as in oocytes with aberrant cytoskeletons and different cytoplasmic streaming conditions. We did so by combining particle image velocimetry (PIV) with Differential Dynamic Microscopy (DDM)^17,18^. In contrast to PIV, DDM probes the sample dynamics not in the direct space, but in the Fourier domain, with the bonus of being able to analyze densely distributed objects, whose size is well below the diffraction limit of the microscope, and in the presence of substantial amounts of noise. DDM can be employed with different imaging mechanisms, providing information on various labeled and unlabeled dynamic structures. These properties are used here for the first time to characterize the crowded interior of a living cell. This is done by combining confocal imaging of labeled F-actin, with simultaneous Differential Interference Contrast (DIC) imaging of unlabeled intracellular vesicles.

Using chemical treatment and genetic manipulation we can show that the dynamics of vesicles result from two contributes: a persistent ballistic motion due to cytoplasmic flows and a diffusion process of active nature. Surprisingly, we found that a similar combination of ballistic and diffusive movements also captures the motility of F-actin, showing a strong correlation between the motion of the actin network and the vesicles. This result provides an important link between the active diffusion of vesicles and the underlying fluctuating non-equilibrium actin network, of which we quantify the overall dynamics by focusing on the diffusive-like component. We discuss our results considering a recently proposed model for the dynamics of a particle embedded in a living cell, where both thermal fluctuations and non-equilibrium activity coexist^19^.

Finally, we demonstrate that, in our system, active diffusion constitutes an ATP-dependent process, with at least two distinct ingredients. On one hand the actin mesh itself seems to be a major source of active diffusion. However, we also find that microtubule based flows enhance active diffusion, and only depletion of both cytoskeletons results in the abrogation of this random motion. As shown before, our data outlines a key role of the dynamic cytoplasmic F-actin in facilitating active diffusion. Importantly, we now demonstrate that microtubules substantially contribute to the diffusive motion of both vesicles and the cytoplasmic actin network as well.

In summary, our work sheds new light on the dynamic interplay between ATP-dependent forces and cytoplasmic mechanics to regulate intracellular motility. We show that: 1) the major ATP-dependent entities responsible for advection and active diffusion are the microtubule and cytoplasmic F-actin networks, and 2) an important dynamic link exists between vesicles and cytoplasmic F-actin motion.

Under a more methodological perspective, we establish DDM as a powerful tool for all biologists interested in problems involving the motion and the rearrangement dynamics of different structures within the cell, as it allows to extract a robust quantitative information even in conditions where more traditional image processing methods fail.

## RESULTS AND DISCUSSION

### Ooplasmic vesicle dynamics consist of persistent and diffusive motion

The asymmetric localization of developmental determinants by microtubule motors (such as kinesin-1) in the *Drosophila* st9 oocyte is a key event for the specification of the body axes of the embryo^20^. In addition, the translocation of cargos by kinesin-1 induces bulk movements of the cytoplasm – called cytoplasmic streaming or flows. These flows can be measured by particle image velocimetry (PIV), using endogenous vesicles as tracer particles^14^. However, while PIV gives an accurate description of flow velocities and topology, it is unsuitable to describe non-persistent, diffusive motion. Therefore, we used Differential Dynamic Microscopy (DDM) to monitor and characterize cytoplasmic movements in more detail (Fig. 1)^18,21^. Analyzing DIC time-lapse movies of living oocytes by DDM (DIC-DDM) unveiled a more complex dynamic behavior, which is not fully captured by PIV. In fact, DIC-DDM analysis shows that ooplasmic vesicles move in a ballistic, persistent, as well as in a random, diffusive manner (Fig. 1a, b).

**Figure 1:**
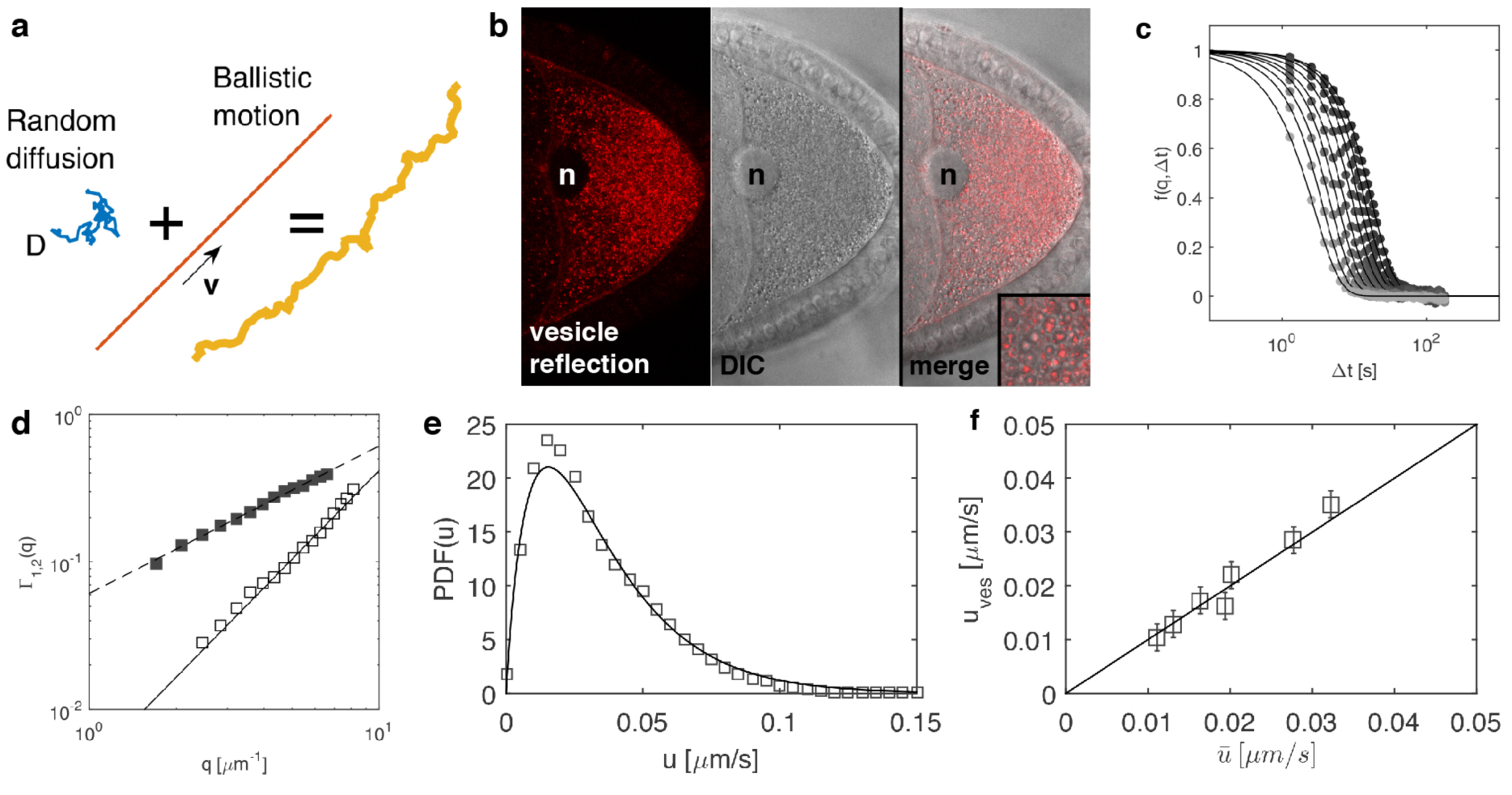
Movement of cytoplasmic vesicles consists of persistent and diffusive motions. (**a**) Diagram, depicting the complex motion of particles in the cytoplasm. Each particle moves by random diffusion, described by a characteristic diffusion coefficient *D* (blue). This random motion can be superimposed to a persistent, ballistic motion, described by a characteristic velocity ***v*** (red). Both together result in the persistent, erratic movement of tracer particles observed in the cell (orange). (**b**) Snapshot of a movie showing reflection of endogenous vesicles in a st9 oocyte, imaged by confocal microscopy (left panel), and the same vesicles imaged by DIC microscopy (middle panel). The merge of both signals reveals that the reflective signal co-localizes with the center of vesicles (right panel, inset). n = nucleus (**c**) Intermediate scattering functions (ISF) *f*(*q*, *Δt*), obtained from DIC-DMM analysis for different wave vectors *q* in the range 1.8 μm^-1^< *q* < 8 μm^-1^. Continuous lines are best fits to Eq. (1). (**d**) Decorrelation rates *Γ*_1_(*q*) (solid black boxes) and *Γ*_2_(*q*) (open boxes) obtained from the fit of the ISF, represented in (c), plotted against the wave vector *q*. *Γ*_1_(*q*), which accounts for the ballistic contribution to the motion of the vesicles, exhibits a linear scaling *Γ*_1_(*q*)=*v*_ves_*q* (dashed line), while *Γ*_1_(*q*), which describes a diffusive-like relaxation process, is well fitted to a quadratic law *Γ*_2_(*q*)=*D*_ves_*q*^2^ (continuous line). 24 (**e**) Probability distribution function (PDF) of the 2D streaming speed in a single cell, as obtained from PIV analysis (open boxes) and as reconstructed from DIC-DDM analysis (solid line). (**f**) Comparison of 2D mean streaming speeds obtained by DIC-DDM (*u*_*ves*_) and PIV 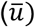 for each cell tested. Both values are in good agreement (solid line represents 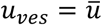), validating DIC-DDM as reliable tool to quantify cytoplasmic flows.

In DDM experiments, the information about the sample dynamics is encoded in the so called intermediate scattering function (ISF) *f*(*q, Δt*), which describes the relaxation of density fluctuations with wave vector *q* as a function of the delay time Δ*t* (Fig. 1c)^18,22^. Initial attempts of fitting our experimental ISFs to the prediction of an advection model, inspired by previous PIV results, failed. Our ISFs were clearly suggesting a more complex, long-time dynamics. However, we obtained a successful description of our ISFs, by using a simple advection-diffusion model, given by

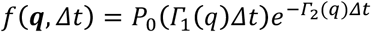

in which, in addition to a directional, ballistic motion with rate *Γ*_1_(*q*), the vesicles are also subjected to a random, diffusive motion with rate *Γ*_2_(*q*). We found a good agreement between our model and the experimental data by considering that each vesicle bears the same diffusivity *D_ves_* (“ves” stands for vesicles), but a different velocity drawn from a prescribed probability distribution function, whose Fourier transform *P*_0_ appears in Eq.1 (see also Materials and Methods). In fact, once the average vesicle speed *v_ves_* is calculated, one has *Γ*_1_(*q*) = *v_ves_q* and *Γ*_2_(*q*) = *D_ves_q*^2^. Fitting the experimental intermediate scattering functions (Fig. 1c) to Eq. (1), confirmed the validity of our model and allowed us to simultaneously determine *Γ_1_* and *Γ_2_* for each *q* (Fig. 1d), both exhibiting the expected ballistic or diffusive scaling, respectively. By repeating this analysis on *N* = 7 cells (with an average of 1500 vesicles per cell contributing to the DIC-DDM signal) we obtained *v_ves_* = 36 ± 15 nm/s and *D_ves_* = (3 ± 1) 10^−3^ μm^2^/s, where in both cases the deviation from the average represents the standard deviation of the population (Table 1).

**Table 1.** Values for diffusive (diffusion coefficient D) and ballistic (velocity v) motions obtained by DIC-DDM (vesicles, ves) or Con-DDM (actin, act) in different genotypes and under different conditions of treatment

| description | $D_{\text{ves}}[\times 10^{-3} \mu\text{m}^2/\text{s}]$ | $D_{\text{act}}[\times 10^{-3} \mu\text{m}^2/\text{s}]$ | $V_{\text{ves}}[\mu\text{m}/\text{s}]$ | $V_{\text{act}}[\mu\text{m}/\text{s}]$ | genotype |
| --- | --- | --- | --- | --- | --- |
| control cells | 3±1 | 6±1.5 | 36±15 | 36±15 | *sqh*-UTRN.GFP |
| no microtubules | 1.4±0.5 | 2.8±1 | nd | nd | *sqh*-UTRN.GFP + colchicine |
| ATP depleted | 0.1±0.5 | 0.14±0.8 | nd | nd | *sqh*-UTRN.GFP + NaN<sub>3</sub>, 2-D deoxyglucose |
| no cytoplasmic F-actin | nd | -- | 150±70 | -- | *spire*<sup>[1]</sup>/Df(2L)Exel6046; *sqh*-UTRN.GFP |
| no cytoplasmic F-actin,<br>no microtubules | 0.6±0.2 | -- | nd | -- | *spire*<sup>[1]</sup>/Df(2L)Exel6046; *sqh*-UTRN.GFP<br>+ colchicine |
| more cytoplasmic F-actin | 1.4±0.5 | 2.3±0.8 | <2 | <2 | *nos*-Gal4,*sqh*-UTRN.GFP>Spire.B.tdTomato |

*nd* = not detectable; -- structure not present; all values given are mean values over all cells ± standard deviation

The ratio *r* = 6 *D*_*ves*_/*v*_*ves*_ ≃ 500 nm corresponds to a characteristic length scale that separates two distinct regimes: over distances larger than *r*, advection is the most efficient transport mechanism, while on smaller length scales diffusion prevails. Of note, in our case *r* is roughly of the order of the vesicle radius. Therefore, it is not surprising that PIV, which operates over a coarse-grained grid with a resolution much larger than the size of the tracers, fails to capture the erratic, small scale, diffusive movement. On the other hand, PIV can efficiently measure the persistent, large-scale motion of the vesicles, which can be used to validate our DIC-DDM approach. To this aim, we analyzed the same DIC movies by PIV (Fig. 1e, f). The comparison between DIC-DDM and DIC-PIV shows that both methods reveal the same quantitative description of flows. With DIC-PIV we find a mean vesicle speed of 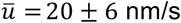 that is in very good agreement with the 2D projection *u*_*ves*_ of the 3D speed *v*_*ves*_ obtained by DIC-DDM, where *u*_*ves*_ = 0.56 *v*_*ves*_ = 20 ± 7 nm/s (Fig. 1e,f, see Materials and Methods for details). Comparable information can also be extracted by PIV analysis of movies, imaging the light reflected by vesicles at 561 nm^14^. We find that each vesicle in the DIC images reflects light at 561 nm from its center (Fig. 1b), and therefore, it is not surprising that they display the same motion (Supplementary Fig. 1). More importantly, the nearly identical velocity values obtained by DIC-PIV and DIC-DDM support the hypothesis that the persistent component of the vesicle’s motion captured by DIC-DDM corresponds to the microtubule-dependent flow previously described by PIV only^14^. However, compared to reflection imaging, DIC has three advantages for general studies on motion of cytoplasmic components. Firstly, the focal plane with DIC is thicker. Secondly, there is no need to use a specific laser line to acquire DIC images, and thus the number of fluorophores that can be combined with DIC is higher. And thirdly, DIC can theoretically be applied to any cell type.

In conclusion, we have shown that DDM can quantitatively separate the persistent, ballistic motion from the random, diffusive-like movement experienced by the same set of vesicles. Importantly, this is obtained through a simple and fully automated procedure, that does not require the accurate localization, tracking and trajectory reconstruction of single tracer particles on which direct-space methods typically rely^23–25^. This makes the results obtained by DDM analysis particularly robust and user-independent, as they are not affected by the selection bias or by the strong dependence on external parameters that are often associated with manual and automated particle tracking. In addition, our method is non-invasive, as no tracer particles need to be injected into the cells, and, furthermore, we do not need to apply any external force to characterize the motion of cytoplasmic components.

Our approach can be of use in a variety of biological problems, involving the characterization of the motion and the restructuring dynamics of different cytoplasmic components, as it provides a detailed and statistically robust description, even in conditions where single particles and trajectories cannot even be resolved or identified.

### The motility of cytoplasmic actin filaments directly correlates with vesicle motion

The cytoplasm constitutes a densely packed environment, containing not only organelles, but also highly dynamic actin filaments. After successfully using DIC-DDM to quantify the motion of vesicles, we applied it to monitor the motility of the cytoplasmic F-actin network traversing the oocyte cytoplasm. This task is made difficult due to the small size of the filaments, their fast and random movement, their crowding, and finally a low signal-to-noise ratio^8,26–28^.

To understand the overall cytoplasmic dynamics in oocytes, we imaged vesicles (DIC) and F-actin simultaneously (Fig. 2a and Supplementary Fig. 2). In st9 oocytes, a three-dimensional F-actin network traverses the entire ooplasm (Fig. 2a-c and Supplementary Fig. 2c)^15^. This actin mesh is formed by the cooperative activity of two nucleators, Spire and the formin Capuccino^15,29,30^. Staining fixed oocytes using the F-actin binding drug phalloidin shows the presence of intertwined filaments, as well as actin rich dots or foci (Fig. 2b,b’ and Supplementary Fig. 2c). However, the dynamic behavior of this structure is completely unknown. Labeling actin in living cells by fluorescent tags has proven to be challenging and yeast formins reject tagged actin monomers^31^ - a fact we could confirm in *Drosophila* oocytes. Fluorophore-tagged Act5C (one out of six actin proteins in flies, and the only one expressed ubiquitously) becomes incorporated into cortical actin structures, but fails to be built into cytoplasmic filaments (Supplementary Fig. 2b). Furthermore, fluorophore-tagged Act5C has a dominant-negative effect, preventing the formation of the actin mesh and inducing fast cytoplasmic flows^32^ (and data not shown). Thus, in order to study the actin mesh in living cells, we used the F-actin binding protein UTRN. GFP, ubiquitously expressed under the *sqh* promotor (Fig. 2c,c’)^33^. UTRN. GFP consists of the calponin homology domain of human Utrophin fused to GFP, strongly binding to actin filaments, but not actin monomers^34^. Based on fixed samples, UTRN. GFP has no effect on the morphology of the F-actin network, or on the timing of its formation and disappearance (Supplementary Fig. 2c). Thus UTRN. GFP constitutes a suitable probe to visualize cytoplasmic F-actin in living oocytes.

**Figure 2:**
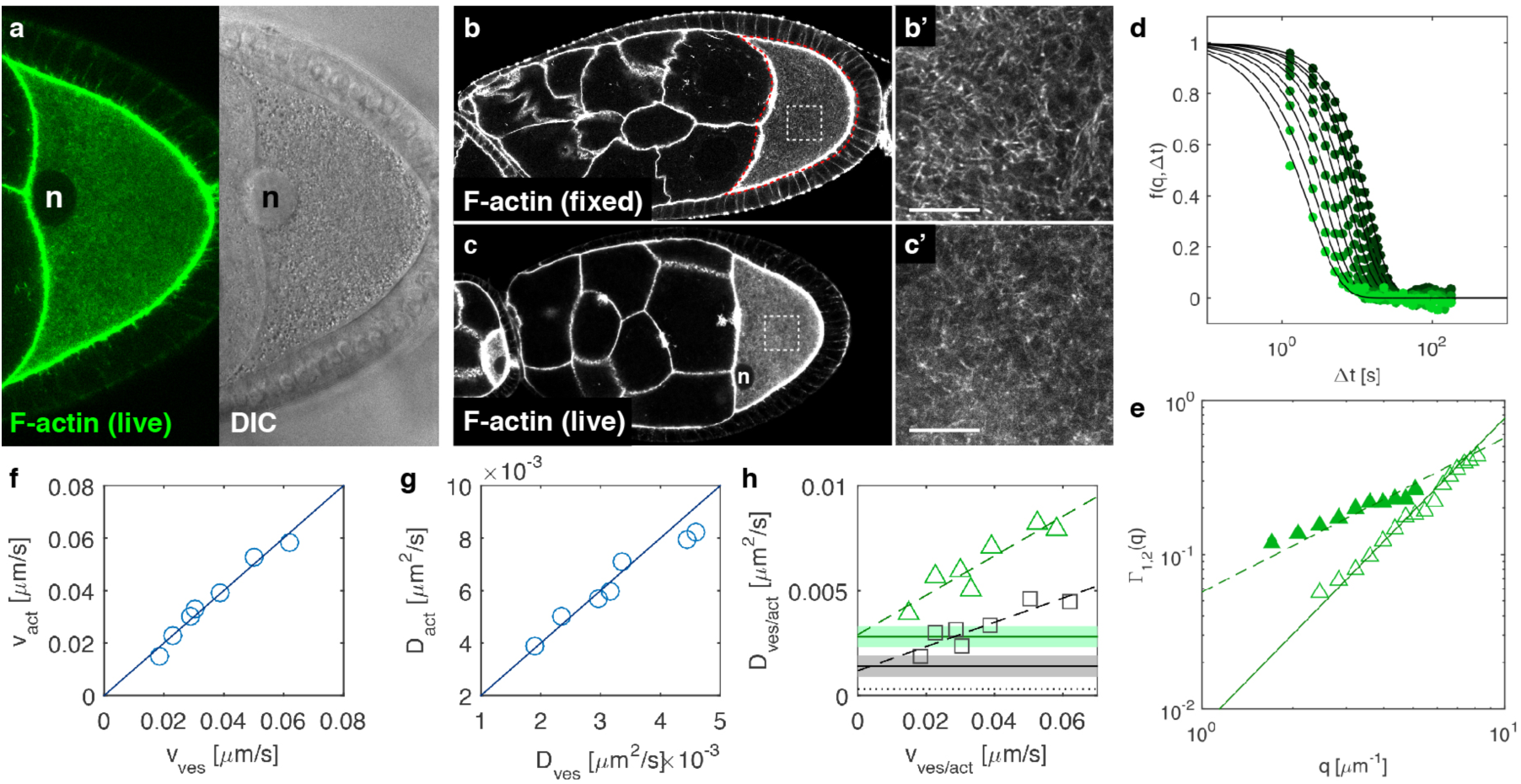
The motion of cytoplasmic F-actin directly correlates with the motion of vesicles. (**a**) Movie snapshot from a cell, expressing the F-actin binding protein UTRN. GFP (left panel, *sqh*-UTRN. GFP). Both the cortex and the cytoplasm of st9 oocytes are enriched in F-actin. DIC image of the same oocyte (right panel). n = nucleus (**b**) Distribution of F-actin (stained with TRITC-phalloidin) in a fixed st9 egg chamber. (b’) High magnification of cytoplasmic actin filaments (white box in b). (**c**) UTRN.GFP expressing living egg chamber. UTRN. GFP labels the same structures as phalloidin in fixed samples (compare to b). (**c’**) High magnification of cytoplasmic actin filaments in a UTRN. GFP expressing living oocyte (white box in C). (**d**) Intermediate scattering functions (ISF)*f*(*q*, *Δt*) obtained from Con-DMM analysis for different wave vectors *q* in the range 2 μm^-1^< *q*<8 μm^-1^. Continuous lines are best fit to Eq. (1). (**e**) Decorrelation rates *Γ*_1_(*q*) (solid triangles) and *Γ*_2_(*q*) (open triangles) obtained from the fit of the ISF, represented in (d). Dashed line constitutes the best fit of *Γ*_1_(*q*) to a linear function *Γ*_1_(*q*) =*v*_*act*_*q*, while the continuous line is obtained from the fit of *Γ*_2_(*q*) to a quadratic function *Γ*_2_(*q*) = *D*_*act*_*q*^2^. (**f**) Mean speeds of F-actin (v_act_), plotted against vesicle mean speeds (v_ves_) with each data point corresponding to one cell. The continuous line represents v_act_ = v_ves_. (**g**) Diffusion coefficients F-actin (D_act_) plotted against diffusion coefficients of vesicles (D_ves_). Each data point corresponding to one cell. The continuous line corresponds to D_act_ = 2 D_ves_. (**h**) D_act_ (green triangles) and D_ves_ (black boxes) as a function of the respective mean speeds vact and v_ves_. While vact and v_ves_ are similar for each cell, D_act_ is consistently higher (~2 fold) compared to D_ves_. Horizontal solid lines represent D_ves, nf_ and D_ves, nf_, obtained from colchicine treated cells, showing no persistent motion (green - F-actin, black – vesicles, also compare to Fig). Dashed areas correspond to mean value ± sd. These values agree remarkably well with the extrapolated behavior for v→0 of the experimental data obtained from control cells (dashed lines). The horizontal dotted line corresponds to the estimated value of the thermal diffusion coefficient D_TH_ of the vesicles, characterizing their spontaneous fluctuation in the absence of any active process. Scale bars represent 10 μm

Visual inspection of the movies revealed that cytoplasmic actin filaments are highly motile and seem to be dragged through the ooplasm in a random, “flowing” manner (Supplementary Movie 1). Those movies also suggest that this F-actin structure does not constitute an interconnected stable meshwork of filaments, but rather resembles a network and of constantly assembling and disassembling filaments, that may intertwine when in close proximity (Supplementary Movie 1). Attempts to quantify the dynamic behavior of the actin network by particle tracking or PIV were unsuccessful, mainly because of the large level of noise and crowding of the structure. Therefore, we combined DDM with confocal imaging (Con-DDM)^35^ to assess its motility. Notably, the same advection-diffusion model used for interpreting the motion of the vesicles is also found to accurately describe the motion of cytoplasmic F-actin (Fig. 2d). Fitting of the q-dependent relaxation rates *Γ*_1_(*q*) =*v*_*act*_*q*, and *Γ*_2_(*q*) =*D*_*act*_*q*^2^ provides an estimate for the characteristic large-scale velocity *v*_*act*_ = 36 ± 15 μm/s and for the effective diffusivity *D*_*act*_ = (6 ± 1.5) 10^−3^ μm^2^/s of the actin filaments (where “act” stands for actin, Fig. 2e, Table 1). Such effective diffusion is in principle a combination of the actual diffusion of the center of mass of actin filaments and of any active process that reshapes the actin network.

The quantities *v*_*ves*_, *D*_*ves*_, *v*_*act*_, *D*_*act*_, display some variability from cell to cell, but exhibit interesting correlations. In fact, comparing the results of F-actin with those on vesicles revealed that the large-scale velocity *v*_*act*_ in each cell compares very well with the vesicle velocity *v*_*ves*_ (Fig. 2f), as *v*_*act*_ ≅ *v*_*ves*_. This result indicates that both F-actin and vesicles move in a persistent manner by advection, most likely driven by microtubule-dependent flows. In addition, we found a remarkable correlation between the diffusion coefficient of vesicles and the effective diffusivity of F-actin, as *D*_*act*_ ≅ 2*D*_*ves*_ (Fig. 2g). This correlation, which will be discussed in more detail below, suggests the existence of an important dynamic link between vesicles and cytoplasmic F-actin motion. This hypothesis is compatible with recent findings in mouse oocytes, where it was found that cytoplasmic actin filaments seem to polymerize from the surface of vesicles^16,36^. However, a comprehensive comparison of all motions (persistent and diffusive) displayed by vesicles (DIC) and the actin network (confocal) was not performed in these studies.

### Microtubule-dependent flow enhances active diffusion

Our results suggest that the diffusive motion of vesicles and actin filaments results from the combination of an intrinsic component and a flow-dependent contribution, which is evident when plotting the diffusion coefficient *D* of either the vesicles or actin as a function of the corresponding velocity *v* (Fig. 2h). The linear dependence of *D*_*ves*_ on *v*_*ves*_ is well captured by the fitting function *D*_*ves*_ = *D*_*ves,0*_ (1 + *a*_*ves*_*v*_*ves*_) with *D*_*ves*,0_ = (1.2 ± 0.5) 10^−3^ μm^2^/s and *a*_*ves*_ = (2 ± 1) 10^−2^s/μm, where *D*_*ves,0*_ is the diffusion coefficient for the intrinsic component 1/*a*_*ves*_= 50 μm/s, the typical velocity above which flows considerably affect diffusion. Notably, the flow-independent diffusion coefficient *D*_*ves,0*_ is significantly larger than the thermal diffusion coefficient *D*_*TH*_= 3.1 10^−4^ μm^2^/s (TH for thermal) that can be estimated for vesicles in the oocyte (average size 1 ± 0.2 μm) and based on previous measurement of the viscosity (η= 1.4 Pa s) for these oocytes (horizontal dotted line in Fig. 2h)^14^. We can thus conclude that *D*_*ves,0*_ describes an active diffusion process, devoid of any flow contributions. Similar results were also found for actin, since the effective diffusivity *D*_*act*_ = *D*_*act,0*_(1 + *a*_*act*_*v*_*act*_) is made of an intrinsic and a flow-dependent contribution, with *D*_*act,0*_ = (3 ±10^−3^ μm^2^/s and *a*_*act*_ = (3 ± 1) 10^−2^ s/μm (Fig. 2h). This suggests that both vesicles and the cytoplasmic F-actin display an active diffusive motion, with a component that depends on cytoplasmic streaming, and a component that does not.

To further assess the influence of flows on cytoplasmic dynamics we first attempted to eliminate kinesin-1, the motor responsible for flows^13,37^. However, the morphology of the actin mesh was massively disturbed in oocytes lacking kinesin, and large aggregations of F-actin were observed instead (Supplementary Fig. 3a,b)^38^. However, it is known that the mesh is present in oocytes that lack microtubules^15^, a fact that we used to test our hypothesis in oocytes obtained from females fed with the microtubule depolymerizing drug colchicine (Fig. 3a,b and Supplementary Fig. 3c, d). In oocytes without microtubules – and consequently without flows - the ballistic movement of cytoplasmic F-actin and vesicles was completely abrogated, while a random diffusive motion was still detected (Fig. 3c and Supplementary Movie 2). Remarkably, the experimental values found in the absence of flows for *D*_*act,nf*_ = (2.8 ± 1) 10^−3^ μm^2^/s and *D*_*ves,nf*_ = (1.4 ± 0.5) 10^−3^ μm^2^/s (where nf stands for “no flow”) are in excellent agreement with the values *D*_*ves,0*_ and *D*_*act,0*_ obtained as extrapolation to zero-flow conditions of the experimental data in control cells, and thereby the presence of flows (Fig. 2h, Table 1). Together, these results demonstrate that microtubule-dependent flows significantly contribute to active diffusion.

**Figure 3:**
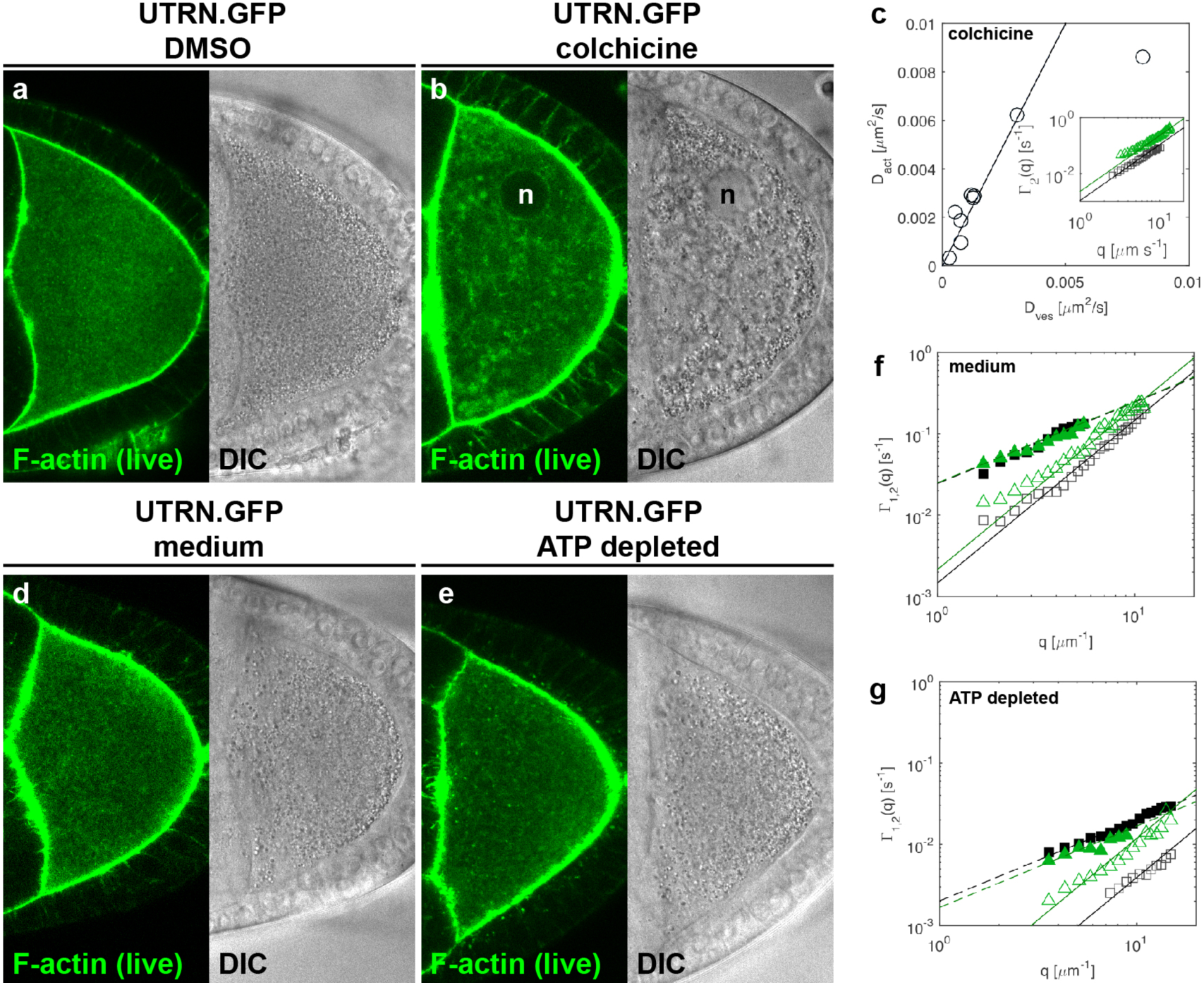
Microtubules and ATP are required for advection and active diffusion. (**a,b**) Persistent motion enhances active diffusion and depends on microtubules. UTRN. GFP (green, left panel) and DIC (right panel) images from living egg chambers, obtained from females fed with DMSO (a) or colchicine (b). The efficiency of colchicine feeding was assured by using only cells that displayed a nucleus (n) detached from the anterior membrane (compared to in Fig. 2a). Depolymerization of microtubules, and consequently the absence of flows, causes a heterogeneous distribution of vesicles. At the same time, prominent actin filaments can be observed traversing the cytoplasm (see also Supplementary Movie 2). (**c**) D_act_ as a function of the diffusion D_ves_. In the absence of persistent flows the diffusivity of both, F-actin and vesicles, is reduced. However, the diffusion coefficients still maintain a linear relation with each other (solid line corresponds to D_act_ = 2D_ves_). Each data point represents one cell. The inset shows the diffusive relaxation rates Γ_2_*(q)* measured for Factin (green triangles) and vesicle (black boxes) in a representative experiment on a single cell. Continuous lines are best fits to the data with a quadratic function. (**d,f**) Cytoplasmic motility is driven by active processes. UTRN. GFP (green, left panel) and DIC (right panel) images from living egg chambers treated with control medium (d) or sodium azide and 2-*D*-deoxyglucose to deplete ATP ((e), see also Supplementary Movie 3)). (f) Γ_1_*(q)* (empty symbols) and Γ_2_(*q*) (solid symbols) measured in a representative control cell (treated with medium). Green triangles correspond to F-actin, while gray squares correspond to vesicles. Continuous and dashed lines are best fits of the data to a quadratic or a linear function, respectively. (**g**) Γ_1_(*q*) and Γ_2_(*q*) obtained from ATP depleted cells plotted against the wave vector ***q***. Color code as in panel (f). Compared to controls, the dynamics of ATP depleted cells is more than one order of magnitude slower.

A final interesting correlation, valid also when microtubules are depolymerized, is that the diffusivity of actin is always faster than the diffusivity of the vesicles, and can be described as *D*_*act*_ ≅ 2*D*_*ves*_ (Fig. 2g,h and Fig. 3c). The active diffusion of vesicles is seemingly locked to the actin mesh, suggesting that the active diffusive motion of the vesicles might originate from an intrinsic activity of the F-actin network. Our results qualitatively agree with a recent model^19^, in which vesicles are caged in the cytoskeleton and the latter act as an actively rearranging harmonic trap for the former. Unfortunately, the rearrangement dynamics of the F-actin network in this model was not explicitly described, as its effect on a tracer particle is described as an effective random force. However, as long as the tracer particle is larger than the actin mesh size, the model predicts that the particle mean square displacement (and so its effective diffusion coefficient *D*_*eff*_) is inversely proportional to its size. In general, we can thus expect in our experiments that a vesicle of size *d*_*ves*_ larger than the actin mesh size *l*_*act*_ will have an effective diffusion coefficient lower than the diffusivity of F-actin (*D*_*eff*_ < *D*_*act*_), with the limiting case being that the size of the vesicle is similar to the mesh size (*d*_*ves*_ ≅ *l*_*act*_), for which we expect an effective diffusion coefficient of the vesicles similar to the effective diffusion coefficient of F-actin (*D*_*eff*_ ≅ *D*_*act*_). In this condition, the vesicle size substantially matches the relevant structural length scale of the network and the particle displacement is expected to closely follow the restructuring dynamics of the trapping mesh. The above arguments are compatible with a simple scaling relation between the diffusivities of F-actin and vesicles: *D*_*eff*_ ≅ *D*_*act*_(*l*_*act*_/*d*_*ves*_). We experimentally found that F-actin diffuses approximately two times faster than the vesicles (*D*_*act*_ ≅ 2*D*_*ves*_) and we can thus expect that the actin mesh size in our experiments is half the vesicle size (*l*_*act*_ ≅ *d*_*ves*_/2).

We evaluated the diameter of vesicle under different experimental conditions. For both control and colchicine treated oocytes we found the same result *d*_*ves*_ = 1.0 ± 0.2 μm. As for the characteristic length-scale *l*_*act*_ for the actin mesh, we calculated the spatial autocorrelation function of the image intensity (Supplementary Fig. 3e). Fitting to an exponentially decaying function provided the characteristic length-scale *l*_*act*_ = 0.42 ± 0.03 μm and *l*_*act*_ = 0.55 ± 0.05 μm, for control and colchicine-treated oocytes, respectively. We thus obtained *l*_*act*_/*d*_*ves*_ = 0.42 ± 0.08 (control) and *l*_*act*_/*d*_*ves*_ = 0.55 ± 0.12 (colchicine) that are both compatible with the expected value *D*_*ves*_/*D*_*act*_ ~ 0.5. These results indicate that the active diffusion of vesicles is due to the non-equilibrium active restructuring dynamics of the underlying actin mesh.

The observed correlation between vesicles and F-actin motility is compatible with recent findings obtained from mouse oocytes, in which vesicles are transported throughout the cell by myosin walking along a F-actin mesh, whose nodes are occupied by vesicles as well^16,36^. An important difference between mammalian and *Drosophila* oocytes is that at the time the actin mesh is present, microtubules are engaged in forming the meiotic spindle in mammals, while in *Drosophila* oocytes, microtubules display an organization similar to a cell in interphase, active in microtubule transport. Despite these differences, our results suggest that the dynamic link between vesicles and actin, and the molecular mechanism responsible for it, are conserved, and survive the presence of microtubule-dependent flows in *Drosophila*. This is further supported by the fact that the murine actin mesh depends on the corporative activity of the actin nucleators Spire and Formin2^39^.

### Cytoplasmic motility is driven by active processes

ATP constitutes the major energy source in cells, and ATP-dependent random force fluctuations have been observed in prokaryotes and eukaroytes^8,10,11^. The observed pattern of cytoplasmic motility in our cells prompted us to experimentally test the active nature of the diffusion-like motion. Therefore, we depleted ATP by treating dissected ovaries with sodium azide and 2-D-deoxyglucose shortly before image acquisition. After acute treatment, the actin mesh remains intact in about half the cells, and ATP depletion leads to an immediate reduction in all cytoplasmic motion. Both, persistent and diffusive behavior of vesicles and actin, are substantially abrogated (Fig. 3d,e and Supplementary Fig. 3f-h and Supplementary Movie 3).

**Figure 4:**
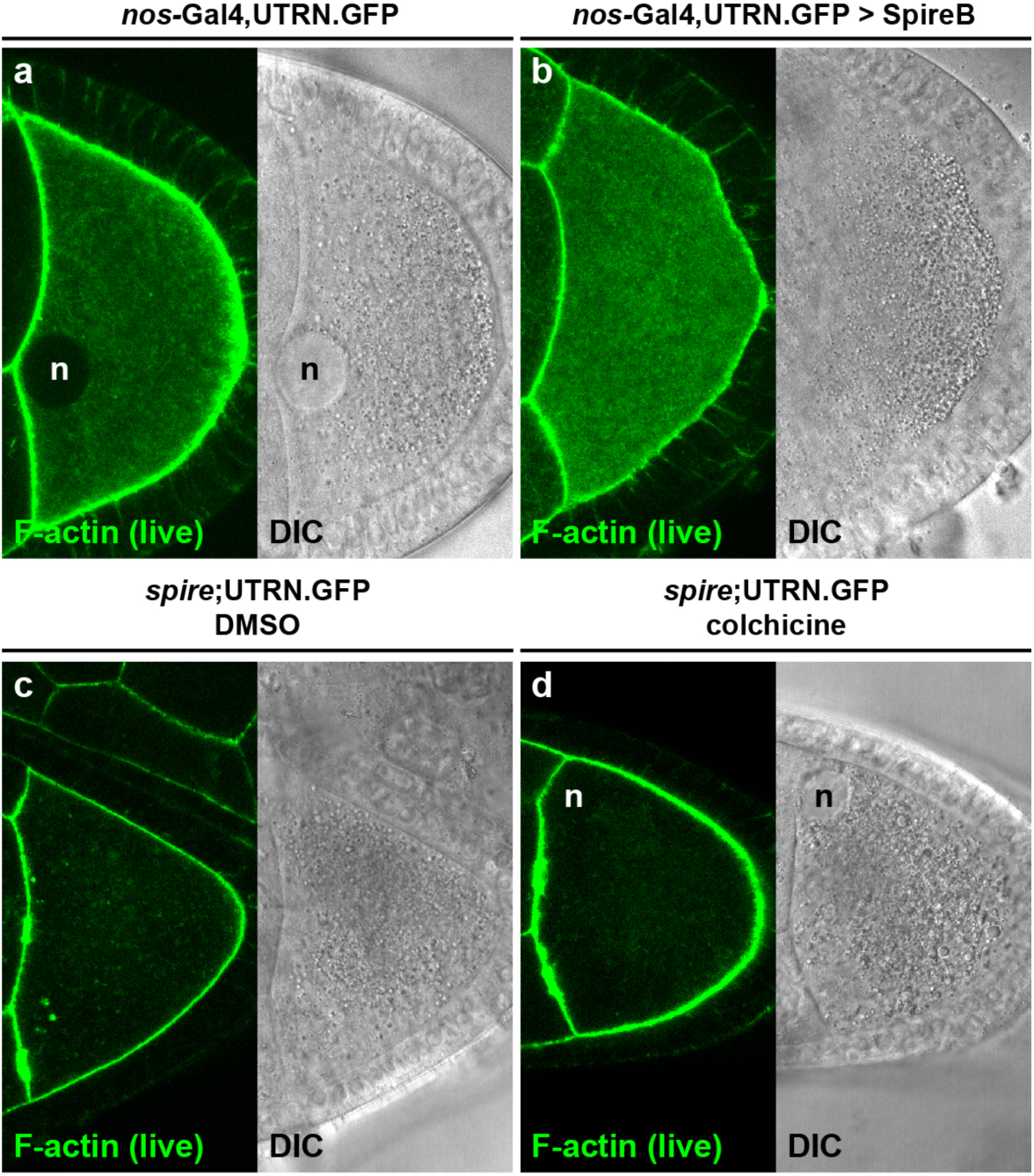
Cytoplasmic F-actin is the major source of active diffusion. (**a**) Living control egg chambers or (**b**) egg chambers over-expressing the actin nucleator SpireB. Overexpression of SpireB causes an increase in the amount of F-actin in the oocyte (see also Supplementary Fig. 4). (**c,d**) *spire* mutant oocytes were incubated in control DMSO-containing medium (c) or in colchicine-containing medium (**d**). As expected, *spire* mutants do not form an actin mesh.

A reliable quantification of the residual dynamics by DDM was difficult, due to presence of a marked plastic deformation of the whole cell (Supplementary Movie 3). Nevertheless, in those ATP-depleted cells, in which this effect was less pronounced, the dynamics of both vesicles and actin could be successfully measured. This was achieved by adopting an advection-diffusion model, like the one described by Eq. (1), where now the advection velocity is assumed to be the same for all particles and, corresponding to the drift velocity, is not significant from a physical point of view. In ATP depleted cells, we find a residual diffusive-like motion, existing on top of the constant drift (Fig. 3f). The average values of the diffusivities associated with vesicles and actin motion are *D*_*ves,ATpd*_ = (1 ± 0.5) 10^−4^ μm^2^/s and *D*_*act,Atpd*_ = (1.4 ± 0.8) 10^−4^ μm^2^/s, respectively (here ATPd stands for “ATP-depleted”). These values are about 30 times smaller than the ones obtained in control cells (Table 1).

Thus, ballistic and diffusive motions displayed by vesicles and cytoplasmic F-actin are strongly ATP-dependent. Since the persistent motion relies entirely on microtubule-based processes (transport and advection), we then asked what the source of the active diffusion is.

### Cytoplasmic F-actin is the major source of active diffusion

F-actin is the source of active diffusion processes in various cell types^8,26 -28^. Therefore, we decided to test whether the cytoplasmic actin network might be driving the diffusive behavior of vesicles in the oocyte. We first tested whether active force fluctuations depend on the level of F-actin in the cell. For this purpose, we studied the motion of both F-actin and vesicles in oocytes that overexpress the actin nucleator Spire. SpireB is the only one, out of four isoforms, to be able to rescue all aspects of *spire* mutants^29^. Nevertheless, the effect of SpireB overexpression in wild type cells has never been studied. Driving SpireB, under the control of the germline specific driver *nanos*-Gal4 (*nos*-Gal4) and the presence of endogenous Spire, results in a dramatic reduction of st9 flows, stratification of the oocyte and complete female sterility (Supplementary Movie 4 and data not shown). The overexpression of SpireB also induces the formation of actin filaments in the cytoplasm of nurse cells (Fig. 4a,b and Supplementary Fig. 4a)^29^. To verify that F-actin concentrations are also higher in the oocyte, we stained for F-actin in fixed cells, using TRITC-labeled phalloidin, and measured the mean fluorescence intensities. We found that the fluorescence intensities in SpireB overexpressing cells are increased by about 1.4-fold compared to controls, showing that the overall amount of actin filaments is increased (Supplementary Fig. 4b). We also measured the mean distance between bright actin filaments as an indicator of the mesh size in the same cells and found no apparent difference between the genotypes (data not shown).

We then analyzed how the overexpression of SpireB and the higher content of actin filaments affect cytoplasmic motility. The most striking effect we found was a strong reduction of cytoplasmic streaming. The ballistic contribution to the overall dynamics was so small that for some cells it could not be reliably measured. Overall, DIC-DDM and Con-DDM analyses provide the following upper bound for the streaming speed: *v*_*ves,SpireOE*_ ≅ *v*_*act,SpireOE*_< 2 nm/s (where SpireOE stands for Spire “over-expression”), a value that is at least one order of magnitude smaller than the typical streaming speed obtained for the control cells. For these Spire over-expressing oocytes, the dynamics are thus substantially diffusive-like, with diffusivities: *D*_*ves,SpireOE*_ = (1.4 ± 0.5) 10^−3^ μm^2^/s and *D*_*act,SpireOE*_ = (2.3 ± 0.8) 10^−3^ μm^2^/s (Table 1). These values are similar to the ones obtained when microtubules are depolymerized, and flows are abrogated. However, in the present case we were not able to reliably study the connection between the dynamics and the spatial correlation properties of actin and vesicles, since SpireB over-expression causes a marked cell-to-cell variability in vesicle size that prevents a meaningful sizing with the available data. Despite this, our data confirm that both flows and active diffusion depend on a well-regulated concentration of cytoplasmic F-actin, and further support a close link between the motion of cytoplasmic F-actin and vesicles.

It is important to note that by over-expressing SpireB in a *spire* mutant background, the mesh is less sensitive to Latrunculin A (LatA) treatment^29^. LatA is a G-actin sequestering substance that not only inhibits *de novo* formation of actin filaments, but also causes the depolymerization of dynamic actin filaments. In wild type oocytes, LatA treatment leads to the destruction of the actin mesh^15^, even within minutes of exposure to the drug^15,40^. However, in oocytes where SpireB expression is driven by an artificial promotor (as in our experiment), LatA treatment remains without any effect^29^. Thus, in SpireB over-expressing oocytes, the aberrant mesh appears to have more, but less dynamic, filaments. This suggests that actin turnover contributes to active diffusion, and it might explain why the diffusive motion of F-actin and vesicles drops in SpireB over-expressing oocytes. Of note, the ectopic expression of Capuccino, a formin jointly working with Spire in forming the actin mesh, also increases the number of actin filaments but does not affect cytoplasmic flows (as measured by PIV)^41^. It is not known why expression of those actin nucleators has a different impact on cytoplasmic flows. However, these observations suggest that a higher concentration of F-actin in the cytoplasm is not enough to substantially inhibit flows.

Finally, we asked how the motion of vesicles changes in oocytes lacking cytoplasmic F-actin. *spire* mutant oocytes do not show any obvious defects in cortical actin organization, but lack the cytoplasmic F-actin network and display premature fast streaming (Fig. 4c,d and Supplementary Movie 4)^15,32,37,42^. DIC-DDM analysis revealed flow velocities in *spire* mutant oocytes to be nearly 5 fold faster than in control oocytes (*v*_*ves*_ = 150 ± 70 nm/s and^29^), making the detection of any superimposed diffusive motion difficult (Table 1 and Supplementary Movie 4). To yet be able to measure diffusivity, we eliminated flows by treating dissected *spire* mutant oocytes with colchicine. In oocytes lacking cytoplasmic actin and microtubules we measured a residual, slow, diffusive-like motion of the vesicles, with an effective diffusion coefficient *D*_*ves,Spire*_ = (6 ± 2) 10^−4^ μm^2^/s (Table 1 and Supplementary Movie 4). This value is about five times smaller than the value measured in control cells and is twice the thermal diffusion coefficient *D*_*TH*_ = 3.1 10^−4^ μm^2^/s, estimated by using the viscosity value obtained in^14^. The diffusivity *D*_*ves,Spire*_ obtained for colchicine treated *spire* mutants is about six times larger than the diffusivity *D*_*ves,ATPd*_=(1±0.5)10^−4^ μm^2^/s measured in ATP depleted cells. This difference might either be due to the presence of microtubule and F-actin independent residual active processes, or by changes in the cytoplasmic viscosity due to the lack of both microtubules and F-actin.

## CONCLUDING REMARKS

In the *Drosophila* oocyte, streaming promotes the mixing and transport of cytoplasmic components^13,43^ and is required for essential events for embryogenesis, such as the localization of mitochondria^4^ and of the developmental determinant *nanos* mRNA^5,45^. Not surprisingly, streaming is more efficient than thermal diffusion in transporting large organelles (e.g. vesicles) over long distances^46^. However, recent studies in cells, which probably lack microtubule-dependent streaming, outlined the important role of active diffusion mechanisms, that at least in mouse oocytes are seemingly dependent on a recently discovered cytoplasmic F-actin cytoskeleton^10,26,27,47^. It is thus worth assessing whether diffusion still plays a role in the presence of streaming and, if that is the case, characterizing the interplay between directed and random motion. In this work, we have developed a robust methodology allowing, for the first time, to separate the persistent motion - due to cytoplasmic flow - from the random motion due to diffusion inside a cell. We have explored how a cytoplasmic F-actin cytoskeleton affects particle dynamics, a question that is largely unknown. In particular, we used Differential Dynamic Microscopy with different contrast mechanisms (DIC and confocal imaging) in combination with genetic and chemical manipulations of the cytoskeletons in the oocyte. Compared to existing techniques, like particle tracking or PIV, the strength of our approach, lies in the ability to quantitatively discriminate between diffusive and persistent motions, without the need of precise tracking (Fig. 5). This enables us to analyze the dynamic behavior of complex, intracellular structures, like the F-actin mesh, which was not possible before. We found that both vesicles and cytoplasmic actin filaments move in a persistent manner by advection, mainly as a result of microtubule-dependent flows. In addition, they display a non-thermal, active diffusion that is dependent on cytoplasmic, but not cortical actin filaments. In fact, we speculate that cytoplasmic F-actin is the major driving force behind active diffusion in the *Drosophila* oocyte. However, we also found that active diffusion is reduced in oocytes without microtubules. This novel finding suggests that microtubules are not only essential for transport and cytoplasmic streaming, but also substantially contribute to active diffusion. This may constitute a difference to murine oocytes, as well as various cultured cells where active fluctuations have been measured^8,28,48^. However, the involvement of microtubules in the cytoplasmic motion of cultured cells was not investigated in detail in these studies, and thus this novel function for microtubules modulating diffusion might be conserved in other systems as well.

**Figure 5:**
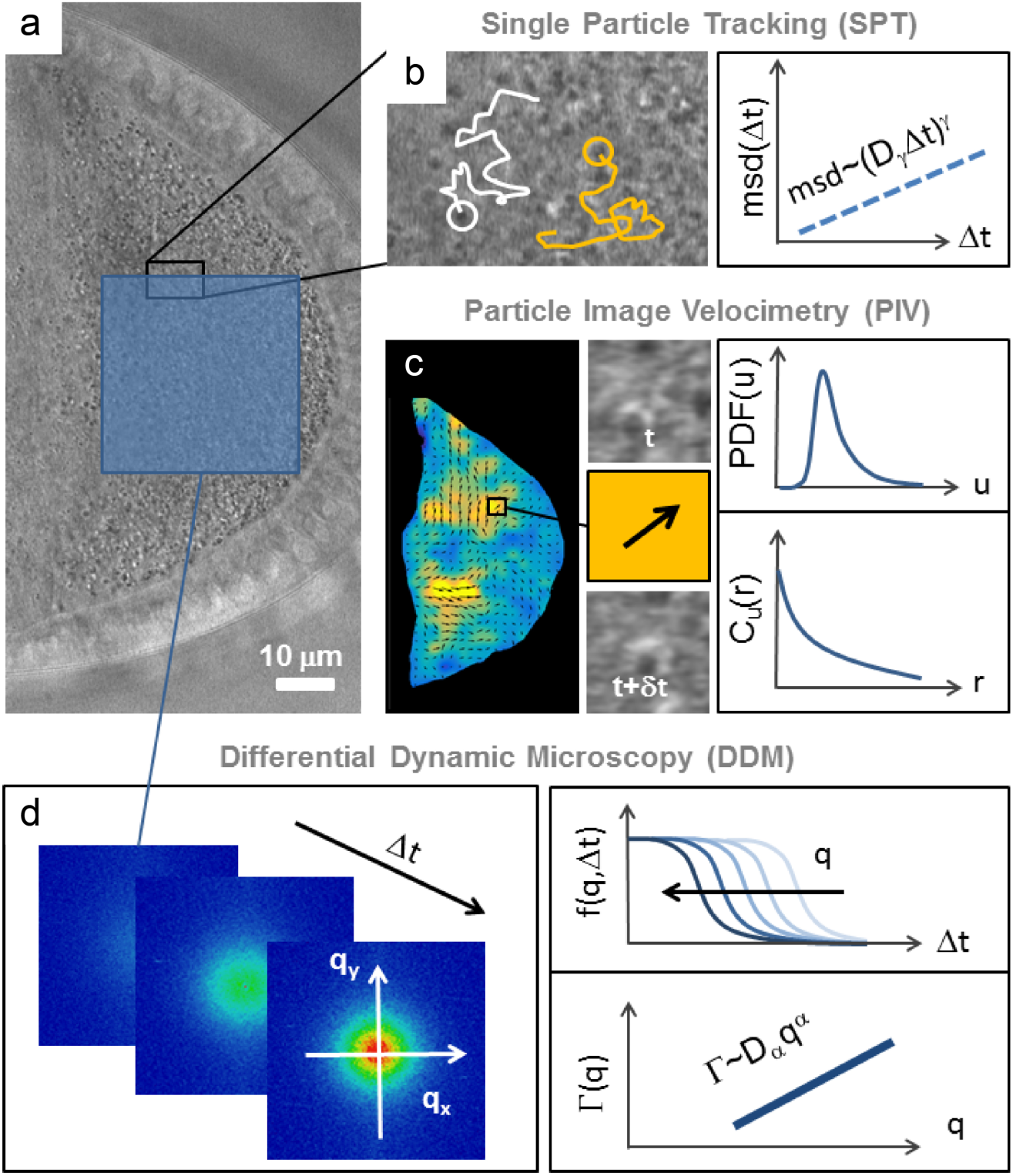
Experimental approaches for the quantification of the intracellular motion. (**a**) Representative snapshot from a movie of a st9 oocyte, imaged by DIC microscopy, where most of the signal originates from the vesicles, densely packed within the ooplasm. (**b**) SPT relies on the identification of specific particles in each frame, and liking their position across different time frames to reconstruct a trajectory (left panel). The typical output of SPT analysis is the mean squared displacement (MSD). The exponent γ, characterizing the dependence of the MSD on the delay time Δt in a given regime, is indicative of the nature of motion (subdiffusive, diffusive, superdiffusive, ballistic etc.), while the prefactor D_γ_ represents a generalized diffusion coefficient. (**c**) PIV enables the reconstruction of a coarse-grained velocity map, where a 2D vector u, representing the local velocity, is associated to each point on a regular grid (left panel). u is determined by estimating the average net displacement across two frames separated by a fixed time delay δt of the particles comprised within a small window centered around the grid point (middle panels). PIV analysis allows the characterization of the statistical properties of flows associated with the ballistic motion of the tracer particles, for example the probability distribution function PDF(u) of speeds or the spatial velocity correlation function C_u_(r) (right panel). (**d**) Compared to SPT and PIV, DDM is not based on the direct space analysis of the images. Instead, DDM quantifies the sample dynamics in the Fourier space, measuring the relaxation of spontaneous density fluctuations occurring at different wave vectors q (left panel). This relaxation process is captured by the so-called intermediate scattering functions f(q, Δt), pictorially represented on semi-logarithmic scales for various values of q. The behavior as a function q of the characteristic relaxation rate *Γ* associated with the decay of f(q, Δ) provides a key information about the nature of the transport mechanism and allows to determine the relevant parameters. The lower left panel pictorially represents the simple and important case where Γ has a power-law dependence on q (log-log scale). This occurs, for example, when the underlying dynamic is diffusive (α=2 and D_α_ is the diffusivity) or ballistic (α=1 and D_α_ coincides with α characteristic speed).

Our study sheds light on the link between the motion of different cytoplasmic components, in particular between large vesicles and cytoplasmic F-actin, which are found to exhibit the same advection-diffusion dynamics. This link is robust upon perturbation of cytoplasmic streaming, and our findings suggest that cytoplasmic F-actin, a dense non-equilibrium fluctuating mesh, is the source of active diffusion for the vesicles. Even though previous cell biological and biophysical observations pointed out that vesicles and the actin network were functionally linked in other cells, our work is the first one to measure and compare the diffusive-like motion of both vesicles and the cytoplasmic F-actin network, finding a quantitative relation between the parameters describing the dynamics of the two structures. In the future, this link will need to be studied in more depth, for instance by performing a more detailed analysis of the correlation between the size of the actin mesh and of the vesicles that we found here. For instance, it would be of interest to study such correlation as a function of the vesicle size.

## MATERIALS AND METHODS

### Fly stocks and genetics

If not stated otherwise, all flies where kept on standard cornmeal agar at room temperature. Fly stocks used in this study where: *w*^*[1118]*^, *w*; *CyO/Sco*; P{*sqh*-UTRN. GFP}/TM6B, *Hu*^*[1]*^*, Tb*^*[1]*34^, *w*; *CyO/Bl*; P{UASp-SpirB.td. Tomato} (gift from M. Quinlan), w;;P{GAL4::VP16-*nos*. UTR} (BL4937), *w*; FRTG13, *Khc*^*[27]*^*\CyO*^49^, *w*; *spir*^*[1]/*^CyO; P{*sqh*-UTRN. GFP}/TM6B, *Hu*^*[1]*^*, Tb*^*[1]*^ (BL5113), *w*; Df(2L)Exel6046/*CyO* (BL7528), w[*]; P{UASp-GFP. Act5C}3 (BL9257) and w[*]; P{UASp-Act5C.mRFP}38 (BL24779).

All UAS/Gal4 crosses were performed at 25˚C, germ line clones where induce using the FLP; FRT *ovoD* system. The stock *w*; *CyO/Sco*; *nos*-Gal4. VP16, *sqh*-UTRN. GFP/TM6B was generated by the recombination of the respective chromosome (this study).

### Actin mesh staining in fixed samples

Ovaries where dissected and fixed in 10% methanol free formaldehyde in PBS, containing 0.1% Tween-20 (PBT0.1), for a maximum of 10 min. The fixed cells were washed 4x in PBT0.1 and stained with 1μM TRITC coupled phalloidin (in PBT0.1, Sigma-Aldrich) over night at 4°C. The stained samples were washed 4x in PBT0.1 and mounted in Vectashield mounting medium (Vectorlabs). Images were acquired on a Leica SP5 inverted confocal microscope, using a 40x/1.3 Oil DIC Plan-Neolfuar or a 100x/1.4 Oil DIC objective. Images were taken within 24h after staining.

### Live imaging of the actin mesh

Ovaries were dissected in a drop of halocarbon oil (Voltalef 10S, VWR) on a glass coverslip and single egg chambers were separated using fine tungsten needles. Images were acquired on a Leica SP5 inverted confocal microscope, using a 40x/1.3 Oil DIC Plan-Neolfuar or a 100x/1.4 Oil DIC objective.

For high-resolution image series, a single plane from the middle of the oocyte was imaged at a scan speed of 100Hz and an image resolution of 1024×1024 pixels (corresponding to one image every 10.4 s).

For DDM analyses images where taken at a scan speed of 400Hz and a resolution of 1024 × 512 pixels, corresponding to one image every 1.29 s. The pinhole diameter was set to a corresponding thickness of the image plain of about 1.39 μm. The cells were illuminated by 488 nm and 561 nm laser light and emission light was collected simultaneously using a hybrid detector at 500-550 nm (GFP), a conventional photon multiplier at 560-650 nm (vesicle auto-fluorescence) and a transmitted light detector with DIC filter set.

### Drug treatment

Depolymerization of microtubules was achieved by either feeding (Fig. 3) or treatment of dissected egg chambers (Fig. 4). For the feeding experiment, 200 μg/ml colchicine were diluted in yeast paste and fed to female flies for 16h at 25˚C. For short-term treatment, flies were fattened overnight, ovaries dissected in dissection medium (1× Schneider’s medium + 2% DMSO) and treated in 20 μg/ml colchicine in dissection medium for 5 min at room temperature (Fig. 4). Ovaries were washed once in a drop of dissection medium and dissected in halocarbon oil. Imaging was performed as described above. ATP was depleted by treating dissected ovaries in 0.4 mM NaN3 and 2 mM 2-Deoxy-D-glucose in dissection medium for 5.5 minutes at room temperature (Fig. 3 and Supplementary Fig. 3f-h). Ovaries were washed in a drop of dissection medium and further dissected and imaged in a drop of halocarbon oil.

### Differential Dynamic Microscopy (DDM)

DDM analysis was performed by using both DIC imaging (DIC-DDM) and confocal imaging (Con-DDM). While Con-DDM was previously demonstrated with densely packed bacteria and colloids ^35^, both techniques are employed here for the first time to probe the cell interior dynamics. Time lapse movies acquired at a fixed frame rate with either imaging mode are treated in the same way. Reciprocal space information is extracted from the analysis of the N intensity frames of the video *i*_*n*_(***x***) = *i*(***x***, *n*δ*t*) acquired at times *n*δ*t* (*n*=1,…, N) by calculating their spatial Fourier transform *I*_*n*_(***q***) = ∫*e*^*j**q**.x*^i_*n*_(*x*)*d*^2^*x*. Here j is the imaginary unit, *δt* is the inverse frame rate of the video acquisition, ***x*** = (*x*, *y*) are the real space coordinates and *q*=(*q*_*x*_, *q*_*y*_) are the reciprocal space coordinates. DDM analysis is based on calculating the *image structure function* defined as

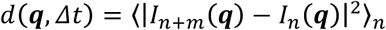

where the average 〈….〉_*n*_ is made over image pairs that are separated by the same time delay Δ*t* = *n*δ*t* over the entire stack. The azimuthal average

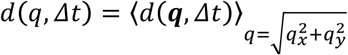

is also typically performed whenever the structure and the dynamics of the sample are isotropic and spatially homogeneous in the image plane, as in the present study. Once the image structure function is calculated, it can be fitted to the following theoretically expected behavior

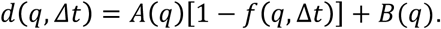

Here *B*(*q*) is a background term due to the detection noise, *A*(*q*) is an amplitude term that contains information about the imaging mode used and the distribution of individual entities in the image, and *f*(*q*, *Δt*) is the so-called *intermediate scattering function*, which encodes the information about the sample dynamics ^17,50^. For instance, diffusive motion with diffusion coefficient *D* is characterized by *f*(*q*, *Δt*)=*e*^−*Dq*^2^*Δt*^ and ballistic motion with constant velocity ***v***_0_ is described by *f*(***q**, Δt*) = *e*^*−jv*_0_·***q**Δt*^ ^18^

A case of interest for the present study is the one of a collection of particles moving via a combination of ballistic and Brownian motion, the first being characterized by a constant velocity *v* drawn from a prescribed distribution *p*(***v***), the second by a diffusion coefficient *D*. In this case the intermediate scattering function reads

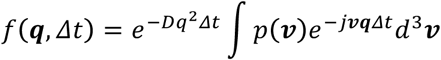

where *p* is the three-dimensional Fourier transform of *p*. If the velocity distribution is isotropic, it can be written in terms of the speed distribution *p*_*s*,3*D*_(*v*)

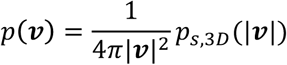

that we assume to be in the form of a Schulz distribution

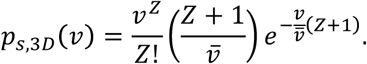

Here *Z* is a shape parameter (set to 2 in our case) and *v* is the average speed:

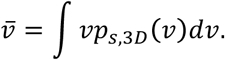

In order to make explicit the dependence on the average speed in the intermediate scattering function one can also consider the rescaled velocity distribution *p*_0_, defined as: 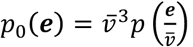. In terms of its Fourier transform *P*_0_ the intermediate scattering function can be written

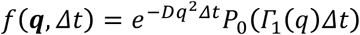

where 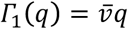 is *q*-dependent decorrelation rate.

If a 2D projection of the motion is considered, the speed distribution reads

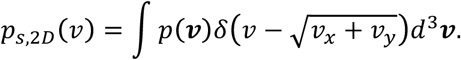

where δ is the Dirac delta function. This is the quantity that is typically measured with PIV. It is important to note that, in general, the mean value of the 2D projected velocity field:

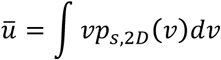

does not coincide with 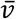. For example, in our case (Schulz distribution with *Z*=2), it can be shown that 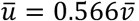.

The choice of the Schulz distribution is motivated by the simplicity of its analytical form, that makes the fitting procedure of the experimental intermediate scattering function particularly simple and robust, and it is supported by the good agreement with the 2D speed distribution measured with PIV (Fig. 1d).

The accessible *q*-range for DDM analysis is given, in principle by [2*π*/*L*, π/*a*] where *L* is the size of the considered region of interest and a is the effective pixel size (typically *L* = 33 μm, *a* = 0.13 μm). In practice, this range can be significantly reduced. This can be due, at low *q*, to the presence of very slow dynamics that cannot be fully captured during the finite duration of the image acquisition and, at high *q*, mainly to the loss of signal due to the microscope transfer function (that depress higher spatial frequencies in the image) and to the presence of dynamics that are too fast to be sampled with the experimental frame rate. Overall, in most of the experiments presented in this work, the effective q-range roughly corresponds to [2, 20] μm^−1^. In the direct space, this corresponds to considering density fluctuations on length scales comprised approximately between 0.3 μm and 3 μm. It is worth noting that, when flow is present inside the oocyte, the velocity field exhibits some spatial heterogeneity, especially close to the cell edges. For this reason, we considered only regions of interest far from these edges, which we imaged long enough to guarantee that the average velocity was rather homogeneous across the field of view.

### Particle Image Velocimetry (PIV)

Maps of the instantaneous intracellular velocities were obtained by analyzing time-lapse DIC and reflection confocal microscopy image sequences with a custom PIV software written in MATLAB ^51^. The time interval between consecutive frames considered for the analysis was 2.6 s. The interrogation window was 32X32 pixels (pixel size comprised between 0.101 μm or 0.145 μm), with an overlap of 50% between adjacent windows. Only the region inside the cell perimeter was considered. Speed probability distribution functions PSD(u) for a single cell were obtained as the normalized histogram of the speeds measured on all grid points and in all frames. Typically, about 1.2·10^5^ vectors contribute to the speed statistics for a single cell.

## ACKNOWLEDGEMENTS

We thank Dr. Margot E. Quinlan for comments and for reagents, and M. Wayland for assistance with imaging. We also thank M. Cosentino Lagomarsino and M. Gherardi for critical reading of the manuscript. MD and IMP were supported by the BBSRC, the Department of Zoology (Cambridge), and the University of Cambridge. FG and RC acknowledge funding by the Italian Ministry of Education and Research, Futuro in Ricerca Project ANISOFT (RBFR125H0M) and by Fondazione CARIPLO-Regione Lombardia Project Light for Life (2016-0998).

### AUTHOR CONTRIBUTIONS

MD, FG, IP and RC designed experiments, analyzed the data, discussed results and wrote the manuscript. MD and FG performed experiments.

